# A Practical Resource for Multi-Omics Data Integration in Microbial Systems

**DOI:** 10.1101/2025.11.19.689359

**Authors:** Warasinee Mujchariyakul, Abderrahman Hachani, Timothy P. Stinear, Kim-Anh LêCao, Benjamin P. Howden, Calum J. Walsh, Romain Guérillot

## Abstract

The increasing availability of microbial multi-omics datasets has created new opportunities to explore complex biological systems. However, exploration remains limited by the lack of accessible, reproducible workflows that integrate multiple omics layers and deliver easily interpretable visualisations of functional and pathway-level insights. Here, we present an R-based workflow for integrated analysis and network-based pathway visualisation of microbial multi-omic data. The workflow enables microbiologists to analyse transcriptomic, proteomic, and metabolomic datasets either individually or in combination, apply univariate and multivariate approaches for biomarker discovery, and generate easily interpretable visualisations of functional and pathway-level signatures. Implemented as multi-step R Markdown, it leverages widely-used open-source tools, including *mixOmics* for biomarker identification and omics integration and *clusterProfiler* for pathway and functional enrichment analyses, with a new network-based integration and visualisation. Its flexible design supports a range of experimental structures and facilitates comparisons across strains, omics layers, and conditions, making it suitable for researchers with limited computational expertise. We demonstrate its utility using a publicly available *Streptococcus pyogenes* dataset, revealing both shared and strain-specific functional responses to human serum. This workflow provides a comprehensive and adaptable framework for systematic multi-omics analysis, improving accessibility and reproducibility and facilitating functional interpretation of microbial responses to diverse environments.

**Data summary:** The code for this workflow is available on GitHub (https://github.com/warasinee/Multiomics_Case_Study). Datasets from our previously published study (1) were used to showcase the functionality and practical utility of the workflow. The multi-omics *Streptococcus pyogenes* dataset used in this study is available in the following public repositories: Gene Expression Omnibus (GSE152821; GSE152822; GSE152823; GSE152824, GSE152826), Proteomics Identifications Database (PXD020863), and MetaobLights (MTBLS2324) (1).

**Impact Statement:** High-throughput omics technologies are transforming our understanding of how microbes adapt to diverse environments and cause disease. The integration of diverse omics layers at a systems level, combining transcriptomics, proteomics, and metabolomics data to identify signature molecules, pathways, and their interactions, remains challenging. Here, we present an R-based bioinformatic workflow designed for microbiology research, which connects existing tools and customised functions to streamline data integration and interpretation. The workflow links biomarkers to functional pathways, visualises results in an interactive network context, and is designed for flexibility and reproducibility. This practical resource lowers technical barriers to microbial multi-omics analysis, providing user-friendly access for dataset exploration and integration and supporting interpretation of system-level microbial adaptation in environmental, clinical, and industrial contexts.

## Introduction

Advances in high-throughput technologies have enabled microbiologists to generate diverse omics datasets, capturing gene expression, protein abundance and metabolite levels, within the same system. Combined, these data offer a multi-layered view of microbial physiology and adaptation, providing opportunities to uncover coordinated responses that single-omics analysis cannot capture. This systems-level perspective contrasts with the traditional reductionist approach often used in molecular microbiology (2), which dissects complex systems into individual components and examines their interactions (3). While reductionism remains essential for elucidating molecular mechanisms, it overlooks the emergent behaviours that arise only when the system is considered as a whole. Consequently, multi-omics integration is therefore becoming a cornerstone of systems microbiology, with applications spanning from bacterial metabolism and stress responses to host–microbe interactions and microbiome ecology (1, 4, 5).

Despite the promise of multi-omics, analysing these datasets remains challenging. Current workflows usually take one of two approaches. In early integration, different omics layers (i.e. genes, proteins, and metabolites) are analysed together from the outset using multivariate methods. A widely used example is DIABLO (Data Integration Analysis for Biomarker discovery using Latent variable approaches for Omics studies), which implements supervised multiblock partial least squares discriminant analysis (PLS-DA) to uncover correlated features across datasets in relation to experimental conditions (5). This strategy helps reveal coordinated biological signals across omics layers. In contrast, late integration involves analysing each dataset separately and then combining the results afterwards, often through pathway enrichment approaches. Two common methods are Over-Representation Analysis (ORA), which tests whether certain pathways contain more differentially expressed genes than expected by chance, and Gene Set Enrichment Analysis (GSEA), which evaluates whether predefined sets of genes are enriched at the top or bottom of a ranked list of all genes (6). Both strategies are valuable, but they are rarely implemented side-by-side in a single, reproducible framework. Moreover, many existing pipelines require users to be proficient in multiple programming languages, and results are often scattered across different tools, which complicates reproducibility and makes biological interpretation less straightforward.

To help address these issues, we present an R Markdown-based workflow that integrates established packages (*mixOmics*, *clusterProfiler*) with custom code and function to connect outputs across stages. The workflow supports both univariate and multivariate analyses, implements early and late integration strategies, and links results to pathway-level network visualisation. By situating biomarkers within functional networks, the workflow enables microbiologists to explore conserved and condition-specific responses across strains, omics layers, and experimental conditions. Rather than introducing new algorithms, our contribution is to integrate established methods into a coherent, R-based modular workflow. This resource is designed to lower technical barriers, enhance reproducibility, and support the functional interpretation of multi-omics data, helping microbiologists translate complex datasets into biological insight across diverse microbial systems.

### Existing Methods and Their Integration in the Workflow

Traditional differential expression (DE) analysis remains a cornerstone of omics research, identifying individual genes or molecules that change between conditions. While powerful, this univariate approach considers each feature in isolation and may overlook coordinated patterns. Multivariate methods, such as those implemented in the *mixOmics* framework (7, 8), address this limitation by analysing many features simultaneously. Techniques like principal component analysis (PCA) allow researchers to explore overall data structure, while (sparse) partial least squares (PLS) and PLS-discriminant analysis (PLS-DA) highlight sets of molecules that best explain or discriminate between biological conditions. For multi-omics integration, multiblock PLS-DA (i.e. DIABLO) extends these methods to uncover signatures that are consistent across different omic layers, providing a system-level view of bacterial responses (7, 8). Importantly, univariate and multivariate analyses often yield distinct yet complementary insights that together help explain complex biological systems. The signature molecules identified through these statistical approaches can then be combined with prior knowledge of underlying molecular pathways and functional annotations using pathway enrichment analysis (PEA) such as ORA and GSEA. To further aid interpretation, we developed a pathway integration visualisation that highlights conserved functional responses to reveals conserved functional responses and supports application to a wide range of multi-omics study designs and biological questions. This network-based integration of functional enrichment can be used to compare responses across bacterial strains, or to combine results from different omics layers and multiple tested conditions when focusing on a single strain (**Figure 1**). The workflow facilitates the identification of biomarkers and signature molecules, contextualising them within pathways and functional interactions to support hypothesis generation and guide functional validation.

**Figure 1.**
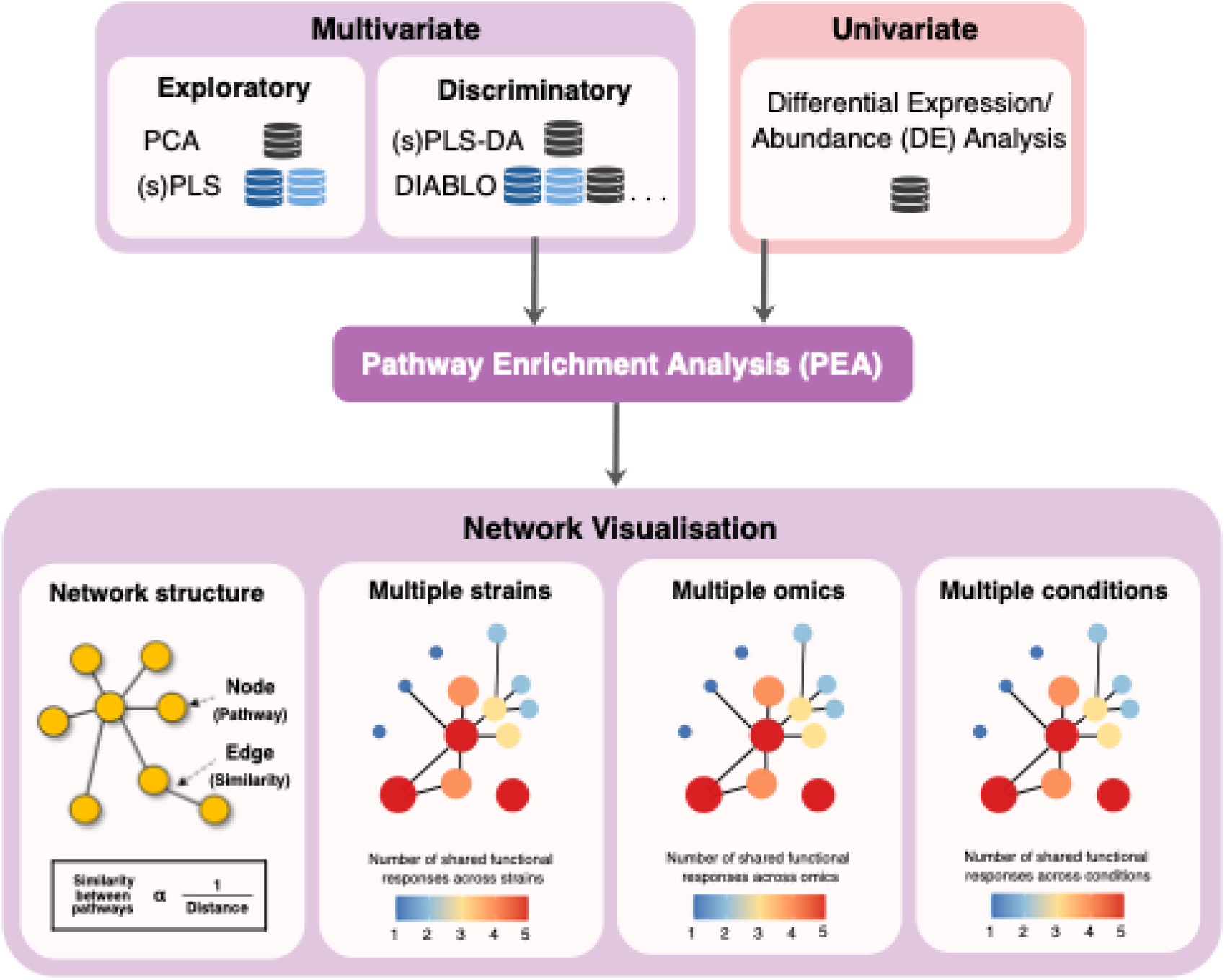
Workflow for Multi-Omics Data Integration and Functional Network Interpretation. Signature molecular features identified from multivariate (i.e., (s)PLS-DA, DIABLO) and univariate (i.e., differential expression analysis) approaches are used as inputs for pathway/functional enrichment analysis (PEA). The resulting pathway-level functional relationships are visualised as networks, with nodes representing pathways or other significantly enriched functional annotations and edges connecting functionally related pathways and functions. Network-based integration of functional enrichment enables the identification of shared functional responses across different strains, omics layers, or multiple tested conditions. The core elements of the workflow are highlighted in purple.

### Comprehensive Overview of the Workflow

This workflow leverages publicly available *R* packages and commonly used software, supplemented with custom *R* functions to connect different analyses’ outputs. The workflow is implemented in *R* Markdown documents that interleave executable code with explanatory text, creating an annotated, step-by-step guide for visualising and interpreting omics results. It supports two major analyses: (i) multi-omics data integration using *mixOmics*, and (ii) pathway enrichment and network analysis (**Figure 2**). Key steps include:

- **Dimensionality reduction and correlation analysis** using PCA and PLS methods in *mixOmics*, enabling exploration of global data structure and relationships across omics layers.
- **Supervised integration and classification** with DIABLO and sPLS-DA from *mixOmics*, which uncover correlated features across datasets and highlight discriminative signatures linked to experimental conditions.
- **Pathway enrichment analysis (PEA)** through *clusterProfiler*, supporting both ORA and GSEA (6), and allowing flexible use of inputs from univariate DE results or multivariate feature rankings.
- **Network-based integration of functional enrichment** of enriched pathways using *Cytoscape* via the *RCy3* interface, with additional interactive visualisation through a dedicated simple shiny app.

**Figure 2.**
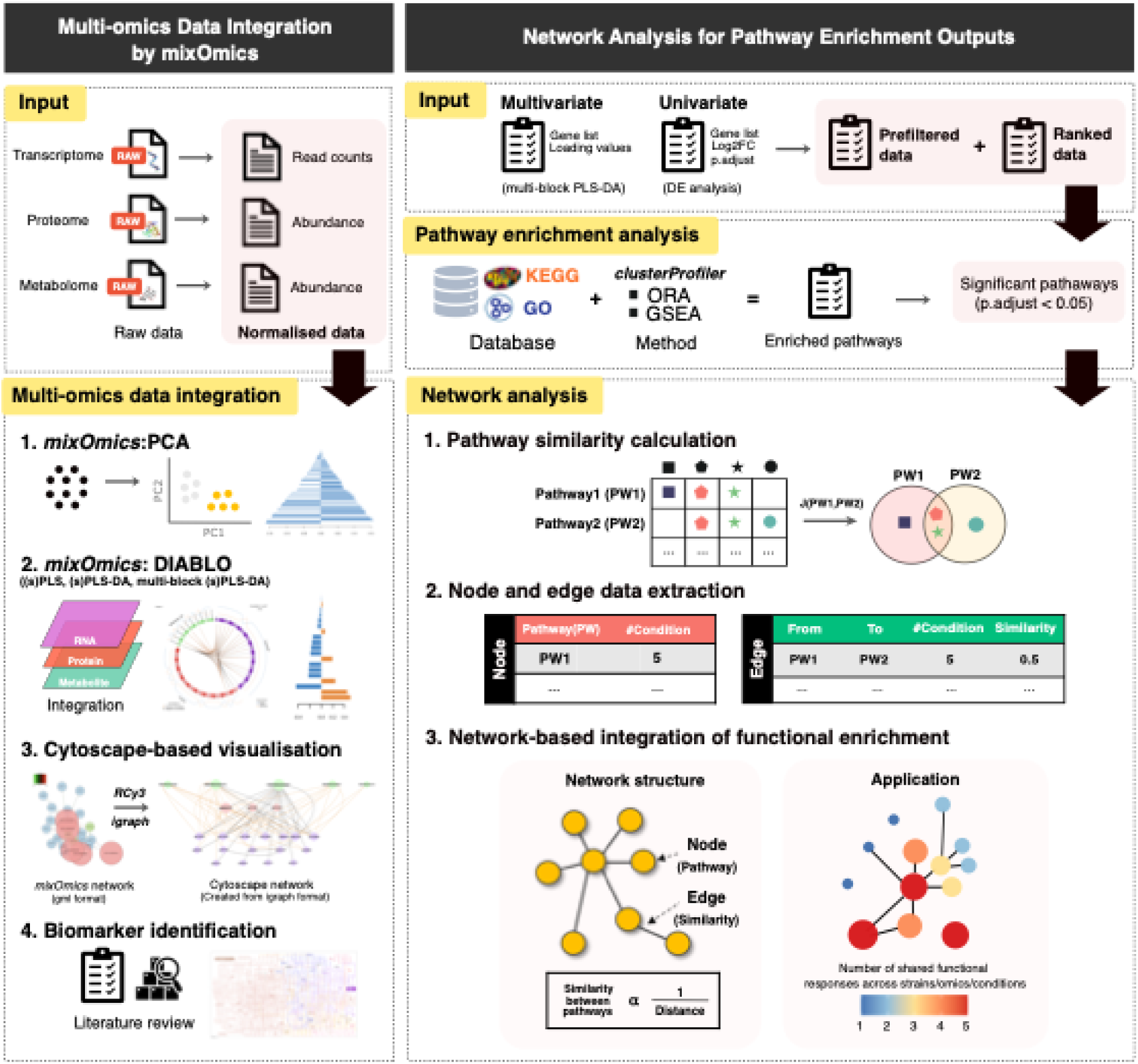
Overview of the integration of multi-omics data, pathway-level network analysis, and interpretation. The workflow combines dimensionality reduction, supervised integration, and network visualisation to identify key biomarkers and functional relationships across omics layers, ultimately leading in network-based analysis of enriched pathways to reveal connected biological processes.

Together, these components situate biomarkers within functional networks, revealing conserved and condition-specific responses across strains, omics layers, and experimental conditions. The workflow thus lowers technical barriers, enhances reproducibility, and supports hypothesis generation by linking statistical outputs directly to biological interpretation.

### Preprocessing, Functional Annotation, and Data Inputs for the Workflow

Preprocessing and quality control are essential prerequisites that lie outside the workflow but ensure reliable downstream analysis. For transcriptomic data, this typically involves read quality assessment, trimming, alignment, and normalization of counts to correct sequencing biases. Proteomic datasets require spectral QC, peptide identification with false discovery rate control, and normalization of quantification values to reduce technical variability. Metabolomic data undergo peak detection and alignment, filtering of outliers, imputation of missing values, and normalization against internal standards to account for instrument drift. These steps collectively minimize noise, remove low-quality signals, and enhance the biological relevance of the inputs carried forward into downstream multi-omics integration and functional analysis. For RNA-seq data, *DESeq2* (9), *edgeR* (10, 11), and *limma-voom* (12) are widely used statistical *R* packages for normalisation and the identification of differentially expressed genes from sequence read counts, with the latter two implemented in the easy-to-use web tool *degust* (13). *MetaboAnalyst* (14), *MZmine 3* (15), and *XCMS* (16) are popular processing pipelines from raw mass spectrometry (MS) data.

Functional gene annotation is essential for linking features to biological pathways and categories. For functional mapping, dedicated microbial annotation pipelines such as *Prokka* (17), *Bakta* (18), *BASys2* (19), *MicrobeAnnotator* (20) and *eggNOG-mapper* (21) are recommended, as they provide per-feature assignments to KO, GO, KEGG, eggNOG, or COG terms, enabling consistent functional annotation across datasets (22). The workflow requires three main inputs: (i) Normalised abundance matrices (features × samples) for each omics layer; (ii) Differential expression results table containing identifiers, experimental metadata, and statistical outputs (e.g., logFC, FDR); and (iii) Functional annotation mappings that link features to pathways or functional terms. Detailed specifications of input file formats, recommended preprocessing steps, and example datasets are provided in the GitHub repository to ensure reproducibility and compatibility with the workflow.

### Case study: Multi-omic characterisation of *Streptococcus pyogenes* response to human serum

The workflow was originally developed to characterise the *Staphylococcus aureus* dataset described in (23). Here, we demonstrate its broader applicability using a multi-omic dataset for *Streptococcus pyogenes*, which includes transcriptomic, proteomic, and metabolomic data (1). In this dataset, samples were collected from five clinically relevant strains grown under two conditions - RPMI or human serum - with six biological replicates per condition, yielding a total of 60 samples.

#### Unsupervised, single-omic analysis

To assess the similarity of bacterial responses to human serum, we performed Principal Component Analysis (PCA) across all omics datasets. The primary source of variation corresponded to the growth condition, indicating distinct global profiles between serum- and RMPI-grown bacteria (**Figure 3A**). Each dataset showed clear separation between conditions, suggesting that *S. pyogenes* undergoes a substantial adaptation to human serum. Shared (conserved) responses across strains accounted for most of the variation, whereas strain-specific effects were comparatively minor (**Figure 3B**). Notably, the transcriptomic and metabolomic profiles of strain SP444 differed from those of other strains, while the proteomic data showed little strain-related separation along the first two principal components (**Figure 3B-C**). To further explore relationships between omics layers, we applied Projection to Latent Structures (PLS), also known as Partial Least Squares. This approach models the covariance between datasets to assess whether shared information exists across layers. Pairwise PLS analyses revealed strong correlations (r > 0.9) between the first latent components of the omics datasets, highlighting a high degree of co-variation and, therefore, a coordinated biological response among the omic layers.

**Figure 3.**
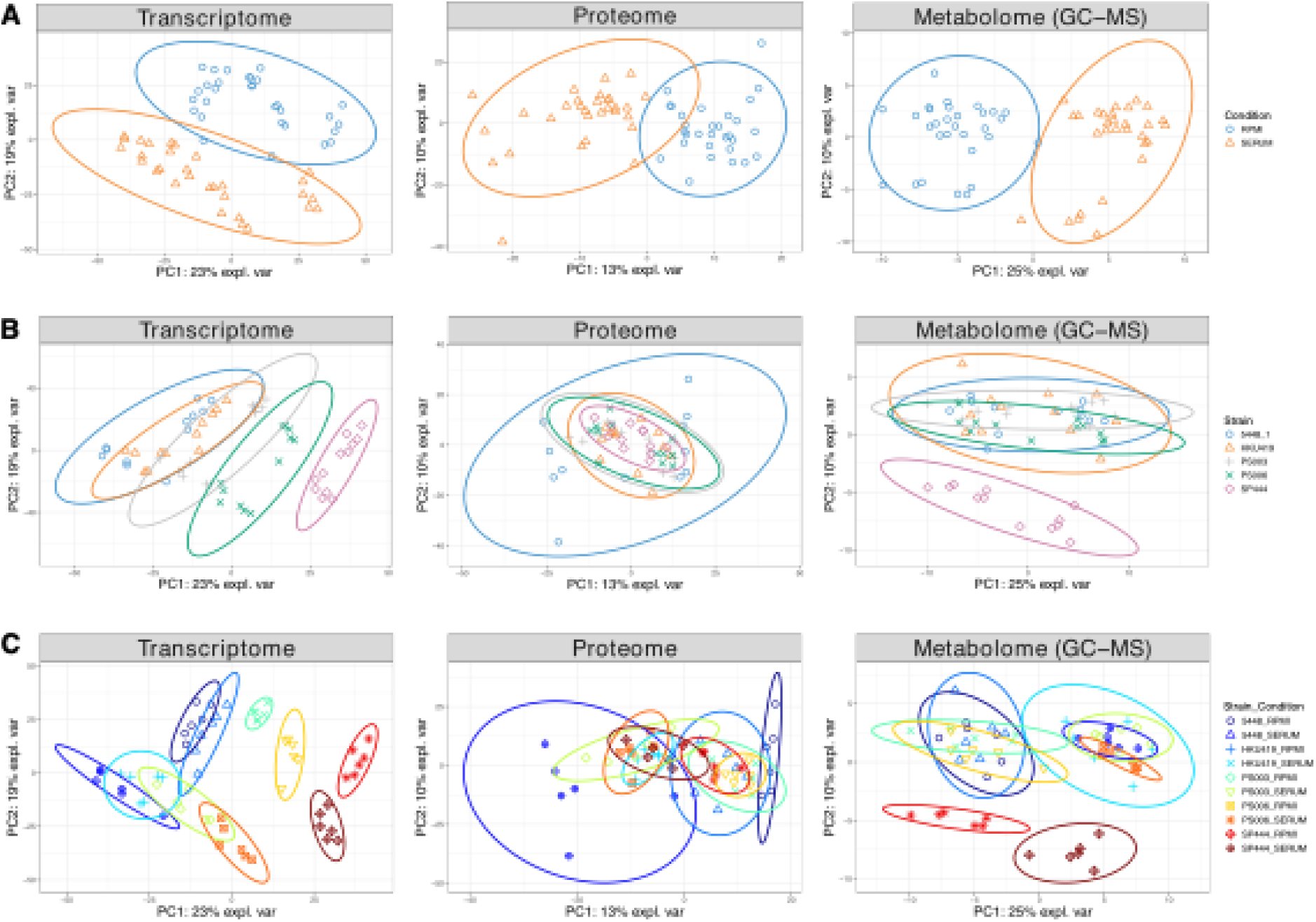
Unsupervised, single-omic analysis of Streptococcus pyogenes response to human serum using unsupervised Principal Component Analysis. Sample plot from PCA illustrating the overall similarity between samples across transcriptomic, proteomic, and metabolomic (GC-MS) datasets. Samples are coloured by growth condition (A) or strain (B) or a combination of strain and condition (C).

#### Supervised Single-Omic Analysis

Next, we applied a supervised classification approach using the sPLS-DA framework to identify features that best discriminate between serum and culture media conditions. This analysis demonstrated the enhanced separation power of the supervised method compared to the unsupervised PCA (**Figure 3A and B**), particularly for the proteomic dataset (**Figure 4A** and **B**), where clustering by condition and strain becomes more distinct. Samples separated clearly by media condition along the first component, while strain-specific variation was primarily captured by the second component.

**Figure 4.**
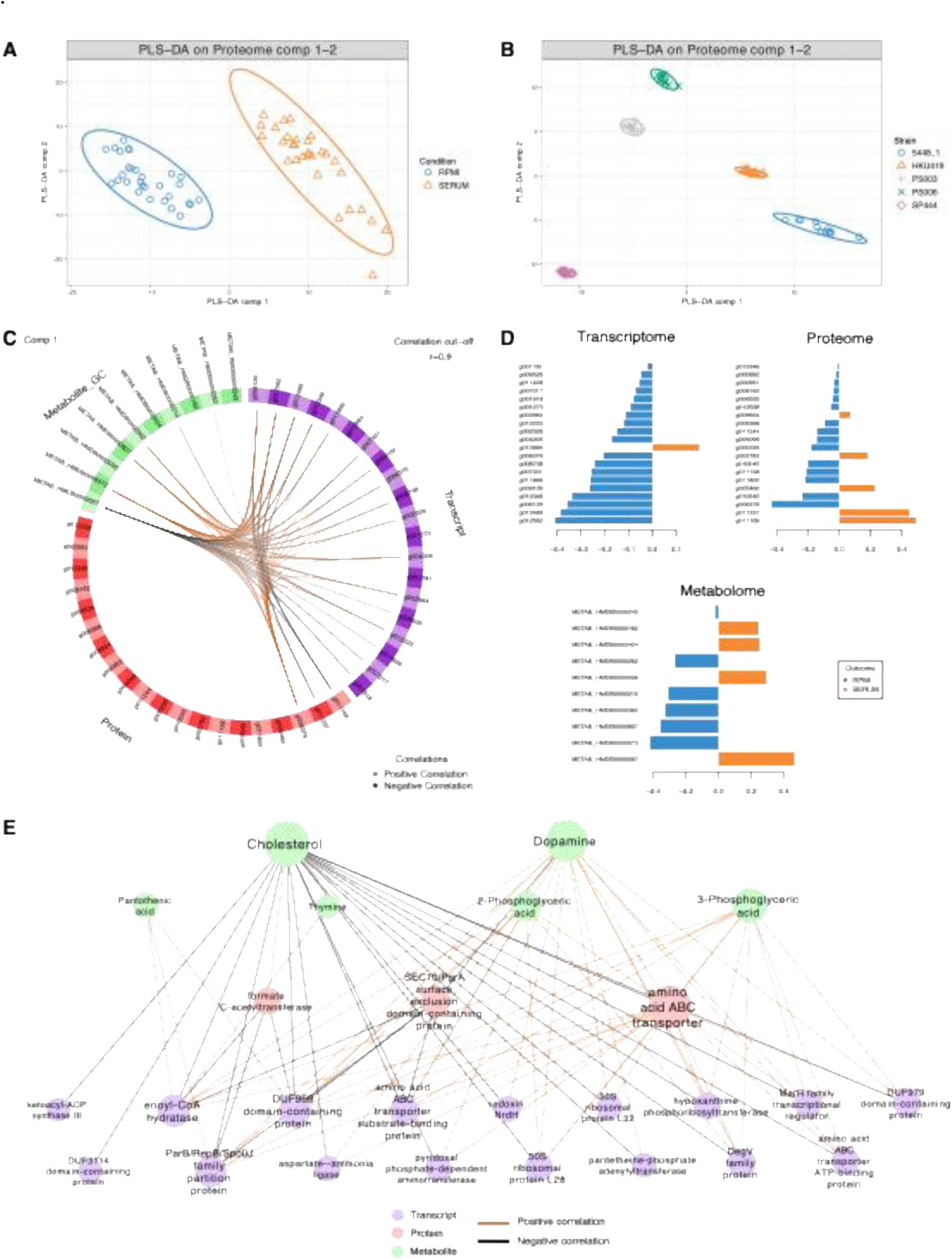
Supervised single- and multi-omic analysis of Streptococcus pyogenes response to human serum. (A-B) Supervised classification using the sPLS-DA framework. Sample plots from sPLS-DA on the proteomic dataset show discrimination of samples by media condition (A) and strain (B). (C-D) Supervised multiblock sPLS-DA (DIABLO) integrating transcriptomic, proteomic, and metabolomic datasets. (C) Circos plot showing pairwise correlations (|r| > 0.9) between features across omics layers. (D) Loading plots illustrating the most discriminatory features along the first component for each dataset. (E) Network visualisation of correlated features, adapted from (C). Edges represent correlations, with edge colours indicating positive (orange) or negative (black) correlations. Node sizes indicate the number of interactions, and node colours represent data type: purple, transcripts; pink, proteins; green, metabolites.

#### Supervised Multi-Omic Analysis

We then employed multiblock (s)PLS-DA (DIABLO) to integrate multiple omics datasets in a supervised framework. The model was trained using labelled data corresponding to the two conditions (RPMI and human serum) and identified highly correlated features across omics layers that maximally discriminate between these conditions (**Table S1**; **Figure 4C–E**). The loading values derived from the DIABLO model (**Table S1; Figure 4D)** reflect the importance of each feature in driving this separation along the first component. Features with larger absolute loading values are the key contributors to the observed differences between conditions. The direction (sign) of these loadings further indicates condition-specific responses – positive loadings correspond to serum-enriched responses, while negative loadings correspond to RPMI-associated features.

#### Pathway Enrichment Analysis

Building on the discriminant and highly correlated variables identified by the DIABLO model, we performed pathway enrichment analysis to explore the biological context of these cross-omic associations. This analysis revealed that carbon metabolism, lipid metabolism, nucleotide metabolism, and defence mechanisms were the pathways most strongly associated with *S. pyogenes* response to serum.

To explore the functional relationships among enriched pathways, we used network-based integration of pathway enrichment results using Jaccard similarity scores to quantify gene overlap between pathways. This approach moves beyond static pathway lists by visualising functional connectivity, revealing both shared (conserved) and strain- or condition-specific responses. Pathway-level networks were generated from three complementary analyses: i) enrichment of discriminant features identified by multi-block PLS-DA, highlighting shared serum responses across strains (**Figure 5A**); ii) enrichment from single-omics differential expression analyses showing both shared and strain-specific responses (**Figure 5B);** and iii) integration of responses across multiple omics layers within a single strain (**Figure 5C**). These networks integrate diverse biological signals into a coherent systems-level view, allowing researchers to identify and prioritize key functions and molecules while providing a foundation for generating testable hypotheses. They revealed major pathways involved in *S. pyogenes* adaptation to serum, including carbon metabolism, ribosomal function, and one-carbon metabolism by folate, as well as pyruvate and butanoate metabolism, which were particularly prominent in strain SP444 based on both transcriptomic and proteomic data.

**Figure 5.**
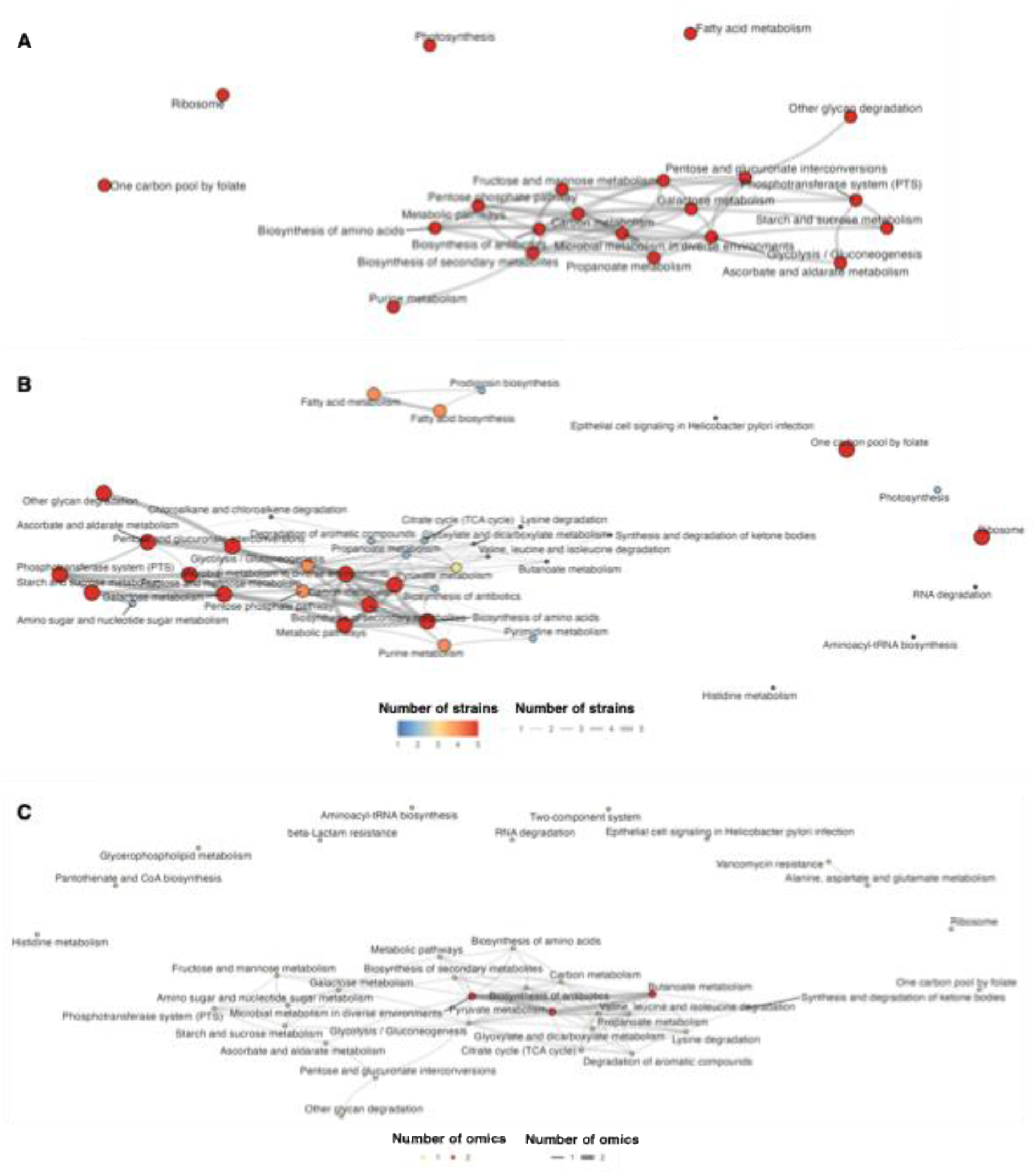
Networks of enriched pathways in Streptococcus pyogenes exposed to human serum. The networks were constructed from the outputs of multi-block PLS-DA (A) and DE analysis (B-C). Nodes represent pathways, and connected edges represent significant pairwise similarity between enriched functional annotations or pathways calculated using Jaccard’s similarity score. The edge length reflects pathway similarity. Node sizes and colours represent the number of strains (A-B) or omics (C) sharing the same enriched pathways. Edge widths indicate the number of strains or omics with interaction between pathways.

Together, this case study uncovered substantial reprogramming of carbon metabolism in *S. pyogenes* when exposed to human serum. Several interconnected pathways showed coordinated shifts, reflecting the bacterium’s metabolic adaptation to the host environment. Many studies have emphasised the importance of specific carbon flow patterns in central carbon metabolism for enhancing pathogen fitness during infection (24, 25) and reprogramming carbon flow has been suggested as a potential therapeutic target (25). Overall, this analysis indicates that *S. pyogenes* likely prioritises host nutrient scavenging over biosynthesis and upregulates fermentation for energy and defence mechanisms.

## Discussion

Rapid advances in high-throughput omics technologies have enabled researchers to capture multiple layers of biological information from a single sample collection. Yet, analyses confined to a single omics layer often fail to fully resolve the complexity of microbial systems, highlighting the need for integrative multi-omics approaches (26). Previous studies have emphasised that the key challenge in multi-omics research is increasingly shifting from data generation to effective downstream analysis, such as data integration, functional interpretation, and biological contextualisation (27–30). Building on established tools, the analysis framework described here delivers a flexible and well-documented approach for microbial multi-omics data integration and visualisation, thereby facilitating biological interpretation.

While this workflow is demonstrated using an existing *Streptococcus pyogenes* multi-omics dataset (1), its structure is generalisable to a wide range of experimental designs and microbial systems. It focuses on downstream data integration, supporting both knowledge-based and data-driven hypothesis formulation, simplifying the use of powerful multivariate and network-based methods for biomarker discovery and pathway-level interpretation within microbial datasets. In particular, the use of a multi-block (s)PSL-DA framework and network integration of pathway enrichment results allows for systematic consideration of multiple biological layers, revealing key molecular targets and pathways relevant to experimental conditions. The network-based analyses applied in our case study capture both shared and strain-specific responses, demonstrating the approach’s capacity to disentangle conserved processes from context-dependent adaptations.

A scale-free network structure was employed because it remains stable even when nodes are removed, reflecting that not all features or pathways are biologically essential under every condition. This design enables the filtering of overly broad categories — for example, general GO terms such as “metabolic process (GO:0008152)” — which can obscure meaningful insights. Focusing instead on more specific terms, such as “glycolytic process (GO:0006096)” and “pentose-phosphate pathway (GO:0006098),” yields more precise and interpretable results in microbial systems.

Looking forward, we anticipate that multi-omics technologies will become routinely used in the scientific community, enhancing our understanding of microbial cellular metabolism and their complex biological systems. The workflow presented here provides an accessible framework to support researchers in exploring, integrating, and interpreting multi-omics data, helping to translate complex datasets into biological insight and testable hypotheses within the expanding landscape of systems microbiology.

## Acknowledgements

We thank colleagues from Howden/Stinear and LêCao labs, particularly Max Bladen, for their technical support and stimulating discussions.

## Author contributions

Conceptualisation, data analysis and visualisation, and writing the original draft and editing (WM); Writing – review and editing (AH, TPS, BPH, KALC). Conceptualisation, methodology, and writing – review and editing (CJW, RG); All authors read and approved the final version of the manuscript.

